# HIF1α-dependent induction of the mitochondrial chaperone TRAP1 regulates bioenergetic adaptations to hypoxia

**DOI:** 10.1101/2020.09.23.309450

**Authors:** Claudio Laquatra, Carlos Sanchez-Martin, Giovanni Minervini, Elisabetta Moroni, Marco Schiavone, Silvio Tosatto, Francesco Argenton, Giorgio Colombo, Paolo Bernardi, Ionica Masgras, Andrea Rasola

## Abstract

The mitochondrial paralog of the Hsp90 chaperone family TRAP1 is often induced in tumors, but the mechanisms controlling its expression, as well as its physiological functions remain poorly understood. Here, we find that TRAP1 is highly expressed in the early stages of Zebrafish development, and its ablation delays embryogenesis while increasing mitochondrial respiration of fish larvae. TRAP1 expression is enhanced by hypoxic conditions both in developing embryos and in cancer models of Zebrafish and mammals. The TRAP1 promoter contains evolutionary conserved hypoxic responsive elements, and HIF1α stabilization increases TRAP1 levels. TRAP1 inhibition by selective compounds or by genetic knock-out maintains a high level of respiration in Zebrafish embryos after exposure to hypoxia.

Our data identify TRAP1 as a primary regulator of mitochondrial bioenergetics in highly proliferating cells following reductions in oxygen tension and HIF1α stabilization.

## Introduction

Changes in metabolic circuities are required by highly proliferating cells during development or tumorigenesis to fuel biosynthetic pathways (Lange *et al*, 2016, Miyazawa & Aulehla, 2018), and central carbon metabolism coordinates a variety of reactions to ensure generation of energetic molecules and anabolic building blocks even in conditions of scarce oxygen availability (Dunwoodie, 2009, Pavlova & Thompson, 2016). To cope with hypoxia, tumor cells can downregulate oxidative phosphorylation (OXPHOS) while increasing their metabolic flux through glycolysis and pentose phosphate pathway (PPP), in a bioenergetic rewiring recalled as aerobic glycolysis or Warburg effect (Warburg, 1956) that allows cells to maintain a high proliferative capacity by providing nucleotides, amino acids and NADPH for anti-oxidant defenses (Vander Heiden & DeBerardinis, 2017). Similar metabolic adaptations are also enacted to boost proliferation during early developmental stages in organisms as diverse as Zebrafish (*Danio rerio*), fruit fly (*Drosophila melanogaster*), African clawed frog (*Xenopus laevis*) and mammals (Agathocleous *et al*, 2012, Miyazawa & Aulehla, 2018, Miyazawa *et al*, 2017). Only later in embryogenesis, the building of the cardiovascular system increases oxygen tension and leads to a concomitant OXPHOS acceleration (Lima *et al*, 2018).

Under limited oxygen availability, the hypoxia inducible factor HIF1α is stabilized and orchestrates a complex transcriptional program that further shapes cell metabolism by inducing glycolysis and repressing OXPHOS (Gordan *et al*, 2007, Lima *et al*, 2018, Zhou *et al*, 2012), promotes cell proliferation and motility and also elicits angiogenesis. Therefore, HIF1α activation is crucial in many settings of neoplastic growth and during embryonic development; indeed, its silencing causes embryonic lethality or developmental defects, depending on the animal model (Compernolle *et al*, 2003, Iyer *et al*, 1998, Kotch *et al*, 1999). We have demonstrated that HIF1α can be stabilized by TRAP1, the mitochondrial paralog of the HSP90 chaperone family, through inhibition of succinate dehydrogenase (SDH) and the subsequent increase in intracellular succinate concentration, which abrogates the priming for proteasomal degradation of HIF1α (Masgras *et al*, 2017a, Sciacovelli *et al*, 2013). TRAP1 induction of HIF1α is pseudohypoxic, as it occurs independently of oxygen availability, and supports a pro-neoplastic metabolic shift toward aerobic glycolysis in a variety of tumor cell types (Masgras *et al*, 2017a). Thus, it constitutes an example of how mitochondria can tune metabolism of proliferating cells, allowing their rapid adaptations to fluctuating environmental conditions. However, it remains poorly investigated the role played by TRAP1 in determining the metabolic changes of cells under physiological conditions, as well as the molecular mechanisms regulating its expression. In this respects, embryonic development, in which cells undergo a fast proliferation rate under varying conditions of oxygen tension (Miyazawa & Aulehla, 2018), represents an interesting field of investigation for TRAP1-dependent regulation of cell bioenergetics.

Zebrafish (*Danio rerio*) is a good model to investigate metabolic features of a developing organism as many of its biochemical pathways are conserved with mammals and its embryonic developmental stages have been extensively characterized (Kari *et al*, 2007, Santoro, 2014). Mitochondrial OXPHOS progressively increases during Zebrafish development through poorly characterized mechanisms (Stackley *et al*, 2011). Here, we report that this metabolic change is accompanied by a parallel decrease in HIF1α and TRAP1 levels. We find that TRAP1 is induced by HIF1α and acts as a major repressor of OXPHOS under hypoxic conditions. These observations apply both to embryonic development and to fish and mammal tumor models.

## Results

### Absence of TRAP1 affects early stages of Zebrafish embryogenesis

We generated Zebrafish TRAP1 knock-out animals using the CRISPR/Cas9 system with a sgRNA targeting TRAP1 exon 6 (**Figures S1A-S1E**). Backcrossing of F_0_ founders with wild-type Zebrafish and subsequent screening of offspring larvae by Heteroduplex Mobility Assay (HMA) allowed the identification of F_1_ germline carriers (**Figure S1B**) with a 13 nucleotide deletion (Δ13) in exon 6 (**Figure S1C**). Homozygous TRAP1 Δ13 fish (Trap1^−/-^) obtained at F_4_ generation (**Figure 1A** and **Figures S1D and S1E**) were used for experimental purposes.

**Figure 1.**
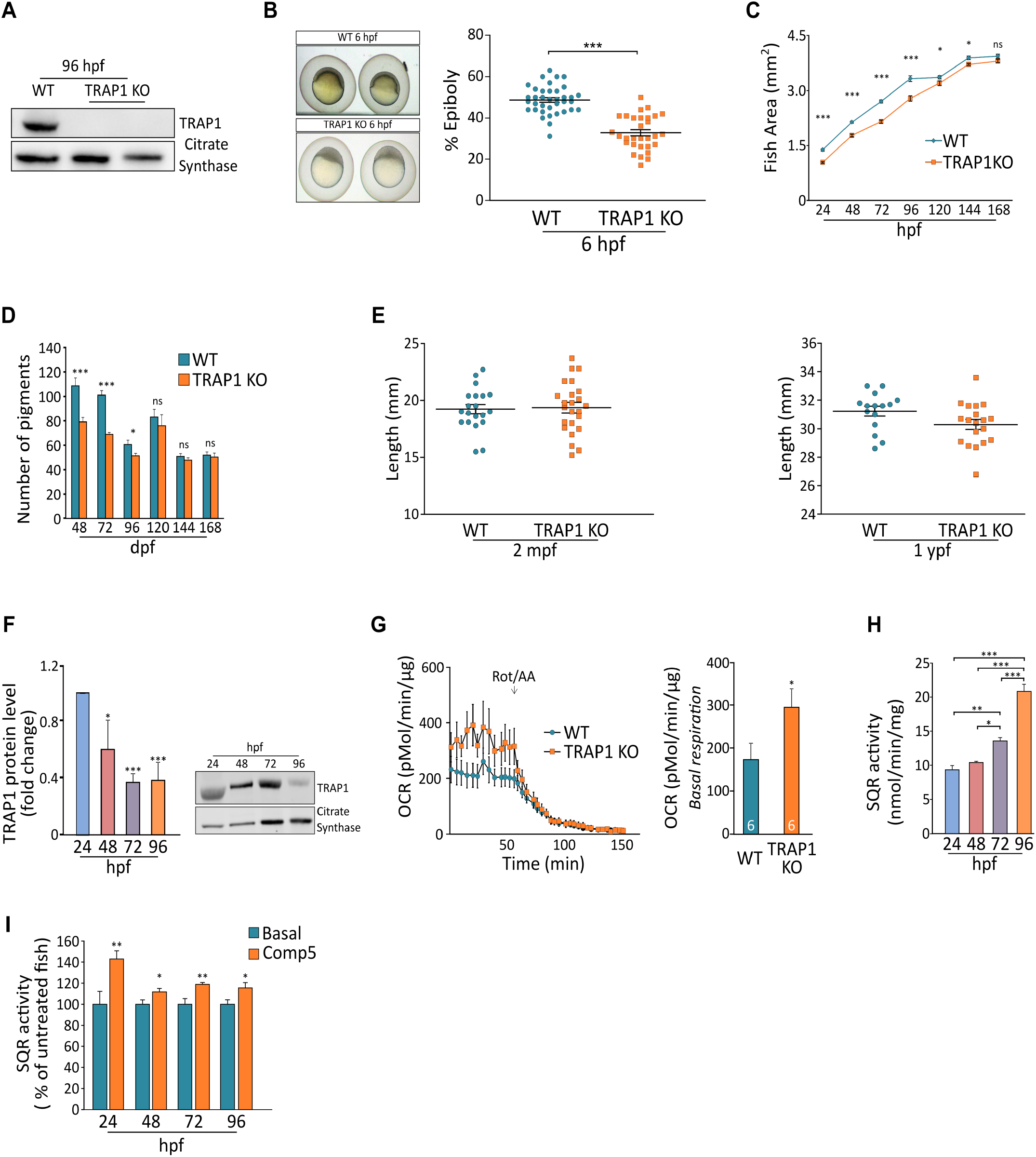
TRAP1 knock-out in Zebrafish affects early embryogenesis stages and mitochondrial bioenergetics. A Western blot analysis of TRAP1 protein level in wild-type and knock-out animals at 96 hpf. The mitochondrial protein citrate synthase was used as a a loading control. B Representative images of developing embryos at 6 hpf and measurements of epiboly area; the yolk was not considered in the analysis. C Kinetics of fish growth during the first 7 days of development. The total fish area, exluding the yolk region, was considered. D Quantification of dorsal pigments from 48 hpf to 7 days post-fertilization (dpf). E Evaluation of wild type and TRAP1 mutant fish length at 2 months post-fertilization (mpf; left) and 1 year post-fertilization (ypf; right). F Analysis of TRAP1 protein expression profile during the first four days of embryogenesis. G Measurement of oxygen consumption rate (OCR) in TRAP1 wild-type and knock-out embryos at 96 hpf. Respiratory complex I and III inhibitors (2 μM rotenone and 5 μM antimycin A, respectively) were added where indicated. Left: representative traces; right: quantification of basal respiration. H, I Succinate-CoQ reductase (SQR) enzymatic activity of succinate dehydrogenase (SDH) in total wild-type embryo lysates. Where indicated, embryos were treated with the TRAP1 inhibitor Compound 5 (100 μM) for 2 hours. In (B-E, I) data are reported as average ± S.E.M. of at least 15 animals and 60 embryos (in I) per condition with an unpaired two-tailed Student’s *t* test. In (G) data are reported as average ± S.E.M. with an unpaired two-tailed Student’s *t* test; the number of animals used for each condition is reported inside column bars. In (H) data are reported as average ± S.E.M. with one-way ANOVA and Bonferroni’s correction of at least 4 independent experiments; asterisks indicate significant differences (*p<0.05, **p < 0.01, ***p < 0.001).

We found that TRAP1 ablation in Zebrafish is compatible with life, but we observed a delay in early fish development. Epiboly, *i.e*. the spreading of cells out of the blastula to form initial sheets of tissues, was markedly retarded at 6 hours post-fertilization (hpf) in TRAP1 knock out larvae (**Figure 1B**). A growth delay of TRAP1 knock-out fish was evident at 24 hpf and maintained during the successive developmental stages until 5 days post-fertilization (dpf) (**Figure 1C**), and the number of pigments was lower in TRAP1 knock-out fish until 96 hpf (**Figure 1D**). These differences were gradually reduced at 5 dpf and completely lost at 6-7 dpf (**Figures 1C and 1D**) and adult fish was not smaller when TRAP1 was absent (**Figure 1E**). These observations are in accord with the changes observed in TRAP1 expression profile during Zebrafish development, as both TRAP1 protein and mRNA declined at 4-5 dpf (**Figure 1F and Figure S1F**), in accord with RNA-sequencing data collected in Expression Atlas database (http://www.ebi.ac.uk/gxa/experiments/E-ERAD-475) that report a reduction in TRAP1 expression during the first 18 developmental stages, which encompass 5 days post-fertilization (**Figure S1F**).

TRAP1 activity could be critical during Zebrafish development in handling oxidative stress linked to the high rate of proliferation coupled to poorly efficient OXPHOS, in accord with the previously reported antioxidant activity of TRAP1 in cancer cell models (Guzzo *et al*, 2014, Masgras *et al*, 2017b). Indeed, we found that TRAP1 knock-out fish displayed a higher level of mitochondrial reactive oxygen species (ROS) than their wild-type counterpart (**Figure S1G**). However, knocking-out TRAP1 is not lethal in Zebrafish and does not hamper its full development, as already observed in mice (Trnka *et al*, 2015).

### TRAP1 regulates mitochondrial bioenergetics during Zebrafish embryogenesis

TRAP1 displays an inhibitory function on OXPHOS in several tumor cell types (Masgras *et al*, 2017b) (Masgras *et al*, 2017a, Yoshida *et al*, 2013), where it down-regulates SDH activity (Sciacovelli *et al*, 2013). Therefore, we wondered whether its absence could affect mitochondrial bioenergetics during early development stages of Zebrafish larvae, when OXPHOS activity and oxygen tension are maintained low (Stackley *et al*, 2011). We observed that abrogating TRAP1 expression enhanced oxygen consumption rate (OCR) in whole living fish embryos (**Figure 1G**). SDH activity displayed a progressive increase within the first 4 days of development, in keeping with the decline in TRAP1 protein levels recorded in the same period (**Figure 1H**). Treatment with the highly specific TRAP1 allosteric inhibitor compound 5 (Sanchez-Martin *et al*, 2020b) increased SDH activity at each developmental stage (**Figure 1I**), strongly connecting TRAP1 activity with SDH inhibition.

### TRAP1 is induced in hypoxic conditions

Our data indicate that TRAP1 levels are elevated during the first days post-fertilization in Zebrafish embryos, when oxygen tension is relatively low. Accordingly, by using a Zebrafish GFP reporter line (Vettori *et al*, 2017), we measured a high transcriptional activity of HIF1α in the early stages of embryonic development, which strongly decreased at 4-5 dpf (**Figure S1H**). These observations suggested the possibility that hypoxia could increase TRAP1 expression. Hence, we exposed 48 hpf Zebrafish larvae to hypoxic conditions for 24 and 48 hours, finding an increase in TRAP1 protein levels that was paralleled by the expected raise in HIF1α expression (**Figure 2A**). Both TRAP1 and HIF1α were already induced after 90/120 minutes of hypoxia exposure in fish embryos both at 96hpf (**Figure 2B**) and at 5 dpf (**Figure 2C**), where basal TRAP1 protein levels were almost undetectable. We also observed a fast induction of TRAP1 following exposure to hypoxic conditions in several human tumor cell types, encompassing MIA PaCa-2 pancreatic cancer cells (**Figure 2D**), U87 glioblastoma cells, BxPC3 pancreatic carcinoma cells and ipNF95.6 plexiform neurofibroma cells (**Figures S2A-S2C**).

**Figure 2.**
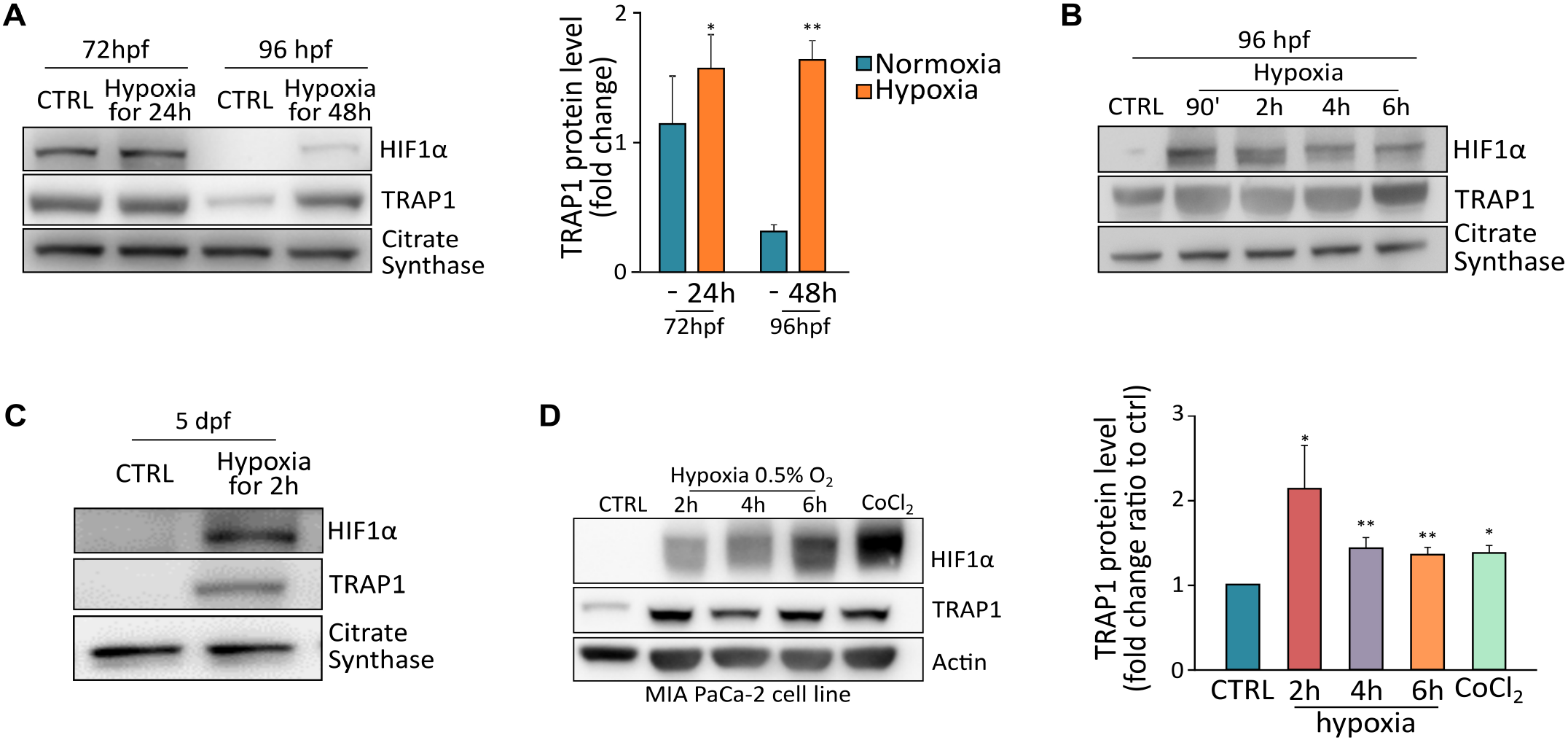
TRAP1 is induced following hypoxia. A Western blot analysis (left) and protein quantification (right) of TRAP1 expression level in embryos exposed to hypoxia (5% O_2_) from the stage of 48 hpf for 24 and 48 hours. B TRAP1 expression profile was analyzed at 96 hpf following short-term hypoxia treatment as indicated. C Western blot analysis of TRAP1 expression in Zebrafish at 5 dpf following short-term hypoxia treatment. D Western blot analysis and quantification of TRAP1 expression in MIA PaCa-2 human pancreatic adenocarcinoma cells following 2-6 hours of hypoxia (0.5% O_2_) or treatment with CoCl_2_ (0.5 mM for 6 hours). The mitochondrial protein citrate synthase and actin were used as loading controls for fish and human cells, respectively. Data are reported as average ± S.E.M. of at least three indipendend experiments with an unpaired two-tailed Student’s *t* test; asterisks indicate significant differences (*p<0.05, **p<0.01, ***p<0.001).

### TRAP1 is regulated in a HIF1α-dependent manner

To evaluate whether TRAP1 induction under hypoxia is caused by its transcriptional upregulation by HIF1α, we searched for hypoxia responsive elements (HREs), the conserved [A/G]CGTG DNA motifs recognized by HIF1 (Kaluz *et al*, 2008), within TRAP1 gene promoter. As the precise extension of the TRAP1 gene promoter is still undetermined in Zebrafish, we explored both a wide genomic region upstream to the Zebrafish TRAP1 locus (*Danio rerio*, Chromosome 3: 9,602,709-9,659,449), including position −5000 bp and +500 bp from the transcription initiation site of the TRAP1 locus (*Danio rerio*, Chromosome 3:9,597,709-9,603,209) (**Figure 3A**). To reduce the probability of identifying misleading motifs, we extended the analysis to the HRE flanking regions assessing a final segment of 33 bp including positions −8 bp up to +20 bp from the canonical HRE (Pescador *et al*, 2005). A preliminary screening identified 94 positive hits homogeneously distributed and showing no preference of localization (**Supplemental Material Table 1**). In order to determine whether any putative HRE may represent an actual HIF1α binding site, we assigned to each HRE a probability value generated using a position-specific frequency matrix (Pescador *et al*, 2005), thus obtaining a final list of 39 highly confident HREs (**Supplemental Material Table 2**). A similar search was repeated on the human TRAP1 promoter region (*Homo sapiens*, Chromosome 16: 3,716,198-3,718,509) finding 43 putative HREs, with 31 sequences presenting a significant probability score (**Supplemental Material Table 3**). Conservation between *Homo sapiens* and *Zebrafish* was then evaluated by sequence alignment that identified four shared sequences containing the motif [A/G]CGTG (**Figure 3B**). These findings are in accord with a HIF1α-dependent regulation of the TRAP1 locus that is evolutionary conserved between Zebrafish and mammals. To reinforce this finding, we extended the analysis by including three supplementary motifs considered to be conserved among hypoxia responsive genes (Schodel *et al*, 2011), finding multiple occurrences of these supplementary hypoxia-related motifs in both human and Zebrafish TRAP1 loci (**Figure 3C**). HREs are the minimal *cis*-regulatory elements required for HIF1α *trans*-activation (Kaluz *et al*, 2008), but hypoxia-dependent gene transcription frequently requires additional transcription factors synergistically acting with HIF1α (Wang *et al*, 1995). GeneCards database (https://www.genecards.org) reported over 30 different transcription factors binding the TRAP1 promoter region. Inspection with STRING database (https://string-db.org/) showed that five of them, *i.e*. SP1, HDAC1, SIN3A, MAX, MYC, are experimentally validated interactors of HIF1α (**Figure S3A**). These findings suggest that HIF1α could mediate TRAP1 transcription acting as a co-transcriptional regulator in tandem with additional transcription factors (**Figure S3B**).

**Figure 3.**
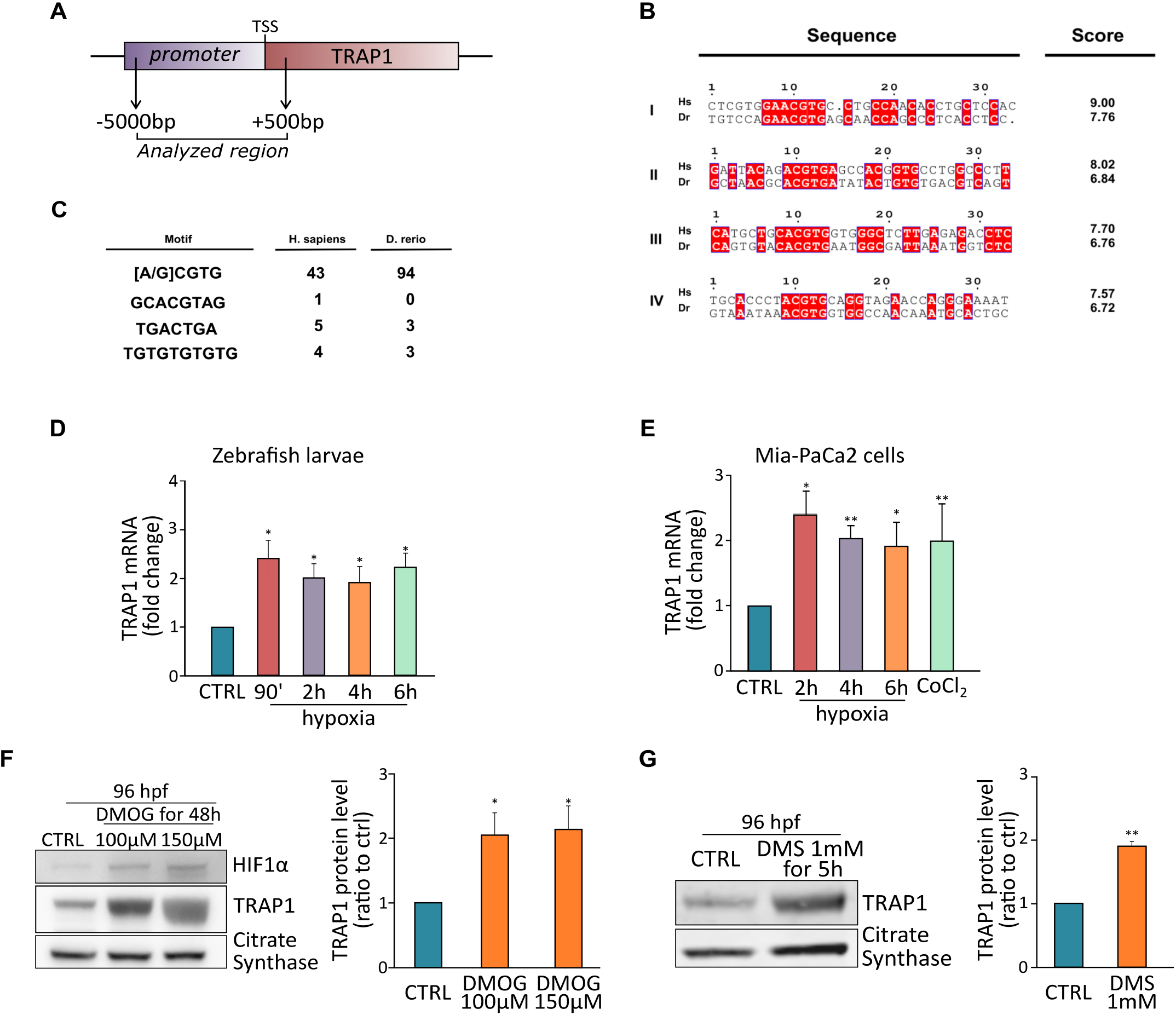
TRAP1 is regulated in a HIF1α-dependent manner. A Schematic representation of the TRAP1 promoter region analyzed that encompasses position - 5000 bp to +500 bp from the transcription initiation site. B List of the four (I to IV) HRE regions more conserved between human (*Homo sapiens*, Hs) and Zebrafish (*Danio rerio*, Dr) TRAP1 promoters. Scoring was calculated using the position-specific matrix. C Relative occurrence of alternative hypoxia-dependent regulative motifs found within the TRAP1 promoter region. D, E Analysis of TRAP1 mRNA expression in wild-type Zebrafish embryos (D) and MIA PaCa-2 human pancreatic adenocarcinoma cells (E) exposed to hypoxia for the indicated times. Values were normalized for expression of rpl13 (D) and actin (E), used as housekeeping genes. F Western blot analysis and protein quantification of TRAP1 expression in embryos at 96 hpf following a 48 hour treatment with 100 μM and 150 μM dimethyloxalyglycine (DMOG). G Western blot analysis and protein quantification of TRAP1 expression in embryos at 96 hpf following a 5 hour treatment with 1 mM dimethylsuccinate (DMS). Data are reported as average ± S.E.M of at least 3 independent experiments experiments with an unpaired two-tailed Student’s *t* test; asterisks indicate significant differences (*p<0.05, **p<0.01, ***p<0.001).

In order to confirm these *in silico* predictions, we exposed to hypoxia both Zebrafish embryos and human pancreatic adenocarcinoma MIA PaCa-2 cells, finding an increase of TRAP1 mRNA levels (**Figures 3D and 3E**). To further assess whether HIF1α stabilization can prompt TRAP1 induction, we inhibited prolyl hydroxylases (PHDs) by using either dimethyloxalilglycine (DMOG) or dimethylsuccinate (DMS), thus preventing HIF1α priming for proteasomal degradation and eliciting pseudo-hypoxic conditions. Treatment of 48 hpf fish embryos with DMOG or DMS strongly induced TRAP1 protein levels (**Figures 3F and 3G**); moreover, knocking-down HIF1α expression almost ablated TRAP1 induction following hypoxia treatment (**Figure S3C**).

Previous reports indicate that HIF1α is stabilized in pancreatic cancer (Hoffmann *et al*, 2008, Schofield & Ratcliffe, 2004, Wilkes *et al*, 2018). We therefore investigated TRAP1 expression in a Zebrafish model of pancreatic adenocarcinoma induced by expressing in the pancreas eGFP-K-Ras^G12D^ with the conditional Gal4/UAS system under the control of the ptf1a promoter (Schiavone *et al*, 2014) (**Figure S4A**). Histological analysis of these fish confirmed the presence of pancreatic adenocarcinoma with mixed acinar and ductal features (**Figure S4B**). In line with published data, we could confirm an activation of HIF1α in pancreatic tumors, as shown by the nuclear accumulation of the HIF1α reporter protein mcherry fused with the nuclear localization signal (nls; **Figure S4A**, lower panel), and an enhancement of HIF1α protein levels in cancer samples with respect to wild-type pancreas (**Figure 4A and Figure S4A**). TRAP1 expression, which was very low in fish pancreas under physiological conditions (**Figure S4C**), was strongly increased both at protein and mRNA levels in K-Ras^G12D^-driven pancreatic adenocarcinomas (**Figures 4A and 4B**). Histochemical investigations of SDH activity on tissue slices showed that SDH activity was strongly reduced in tumor masses compared to normal pancreas, and incubation with the specific TRAP1 inhibitor compound 5 restored the SDH enzymatic activity in samples of pancreatic tumors (**Figure 4C**).

**Figure 4.**
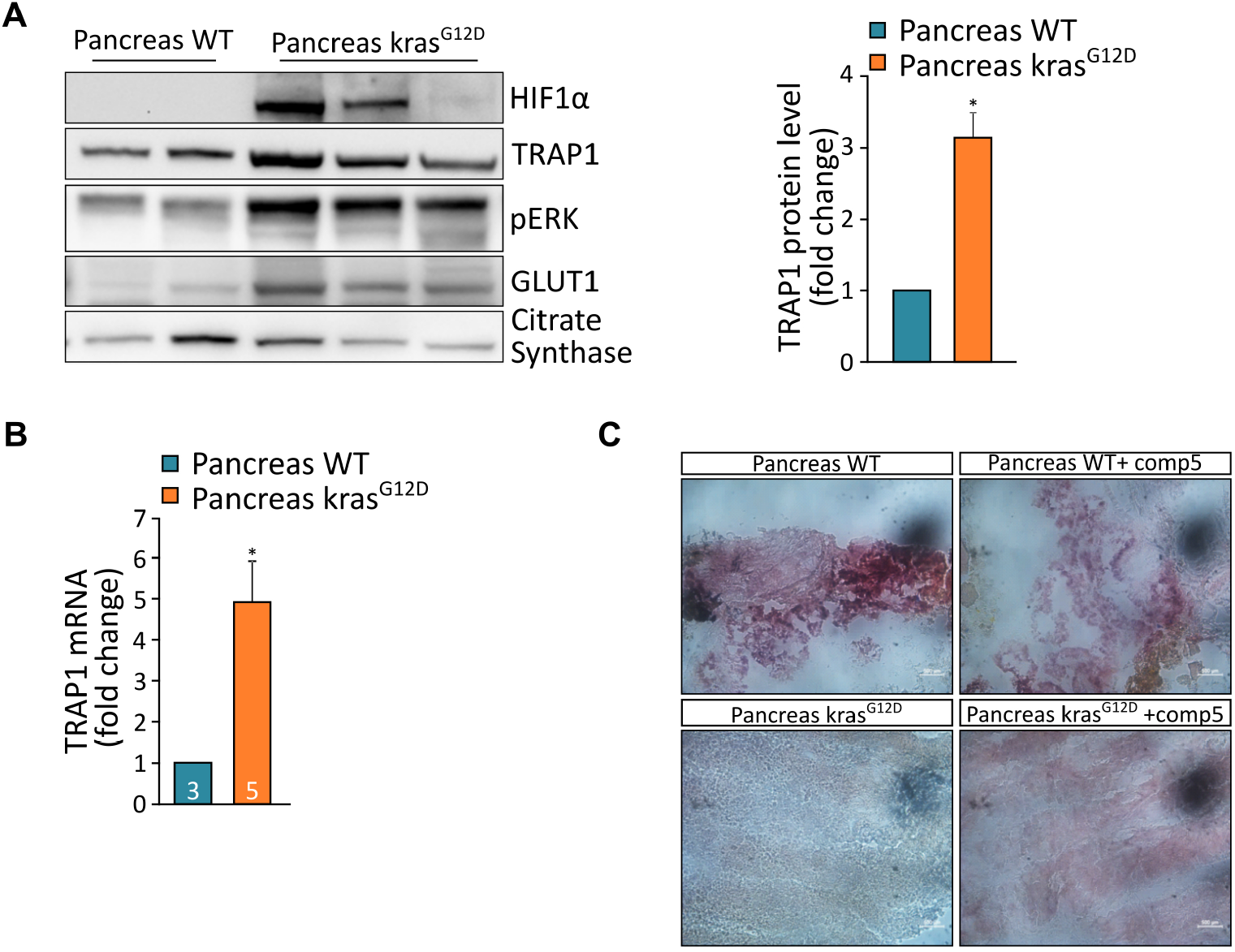
TRAP1 is overexpressed in pancreatic tumors and down-regulates SDH activity. A Western blot analysis and protein quantification of TRAP1 in wild-type pancreas and in pancreatic tumors driven by KRAS^G12D^ expression (1 ypf). Values were normalized for citrate synthase. B Analysis of TRAP1 mRNA expression in wild-type pancreas and in pancreatic tumors at 1 ypf. Values were normalized for expression of rpl13 and actin, used as housekeeping genes. The number of animals used in each condition is reported inside column bars. C Colorimetric assay (pink staining) of succinate dehydrogenase (SDH) activity in tissue slides of wild-type pancreas and pancreatic tumors with/without the TRAP1 specific inhibitor compound 5 (100 μM, 30 minutes pre-incubation). In (A, B), data are reported as average ± S.E.M of at least 4 different animals per condition with an unpaired two-tailed Student’s *t* test; asterisks indicate significant differences (*p<0.05).

Taken together, our data indicate that TRAP1 is induced in a HIF1α-dependent way under both hypoxic and pseudo-hypoxic conditions, and its induction has important consequences on the *in vivo* metabolic rewiring of neoplasms.

### TRAP1 regulates mitochondrial respiration during hypoxia

We then asked whether TRAP1 could contribute to the bioenergetic features of hypoxic cells. Hypoxia slowed down embryo development, an effect exacerbated by the absence of TRAP1 (**Figure S5A**). As expected, a prolonged exposure (48 hours) of Zebrafish embryos to hypoxia lowered both their SDH activity (**Figure 5A**) and OCR (**Figure 5B**). Treatment with the selective TRAP1 allosteric inhibitor compound 5 (**Figure 5A**) or with the non-specific HSP90 family inhibitor 17AAG (**Figure S5B**) rescued SDH activity to levels comparable to normoxic fish, indicating that SDH inhibition in hypoxic Zebrafish larvae was TRAP1-dependent and reversible. Similarly, the reduction in basal OCR observed by placing embryos under hypoxic conditions was completely lost when we abrogated TRAP1 activity, either by compound 5 administration or in TRAP1 knock-out fish (**Figure 5C upper panel and Figure 5D, blue bars**). Remarkably, hypoxia could not decrease basal OCR in TRAP1 knock out embryos, nor the TRAP1-targeting compound 5 affected mitochondrial respiration in normoxic or hypoxic conditions (**Figure 5C lower panel and Figure 5D, orange bars**). Of note, for technical reasons OCR measurements were performed briefly after returning larvae to normoxia, indicating that hypoxia-mediated respiratory inhibition was maintained following the shift to atmospheric oxygen tension, yet it did not occur in the absence of TRAP1 activity. These observations point towards a primary role of TRAP1 in regulating mitochondrial metabolism of Zebrafish embryos exposed to hypoxic conditions.

**Figure 5.**
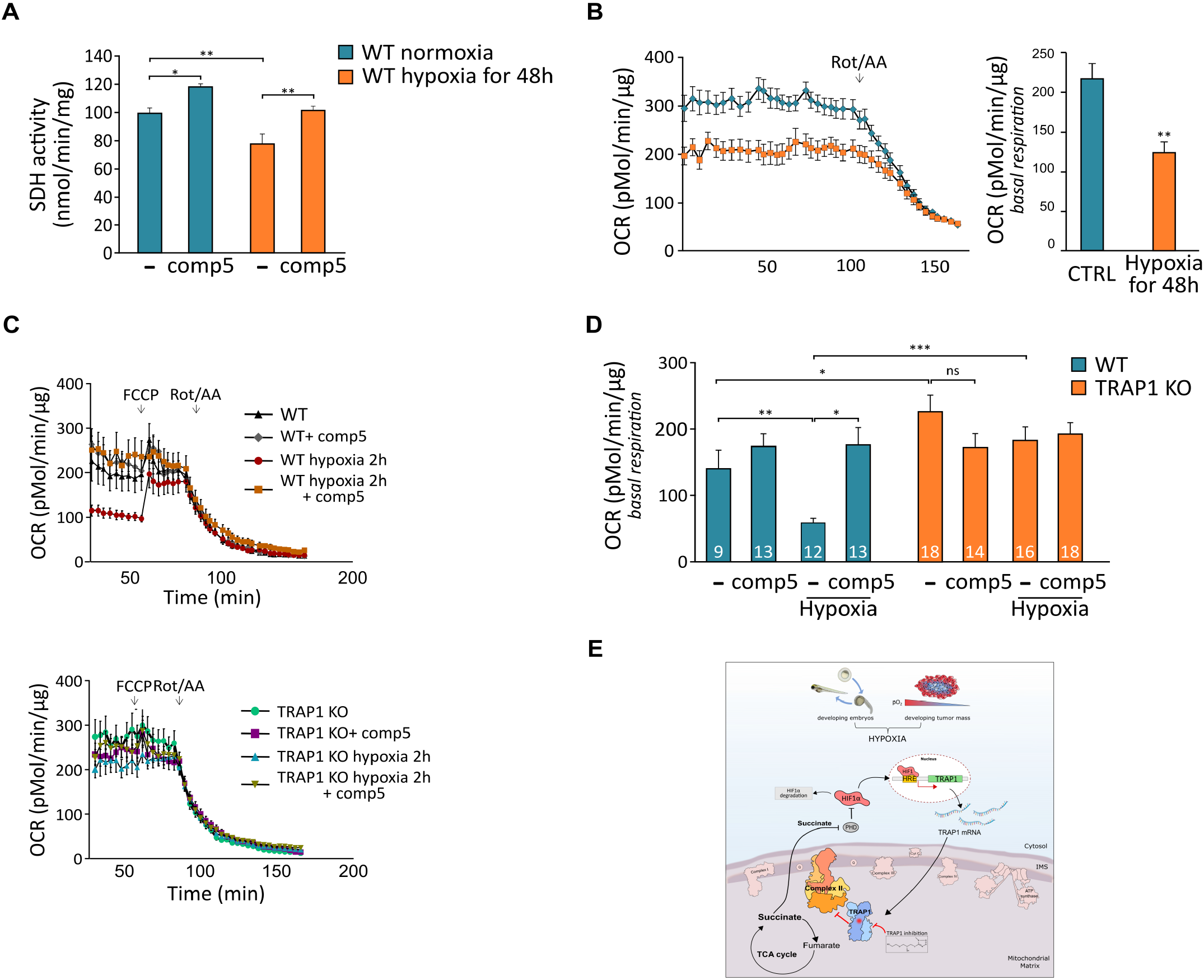
TRAP1 inhibits respiration under hypoxia. A Spectrophotometric measurements of SDH activity on wild-type Zebrafish embryos kept in normoxia or hypoxia (5% O_2_) for 48 hours. Where indicated, embryos were treated with the specific TRAP1 inhibitor compound 5 (100 μM, 2 hours). B Assessment of oxygen consumption rate (OCR) in wild-type embryos at 96 hpf, either untreated (blue line and bar) or exposed to hypoxia for 48 hours (orange line and bar). Respiratory complex I and III inhibitors (2 μM rotenone and 5 μM antimycin A, respectively) were added where indicated. C, D OCR measurements in TRAP1 wild-type and knock-out living Zebrafish embryos, either kept in normoxic condition or after 2 hours of exposure to hypoxia (5% O_2_). Subsequent additions of the proton uncoupler carbonyl cyanide-4-(trifluoromethoxy)phenylhydrazone (FCCP, 0.5 μM) and of the respiratory complex I and III inhibitors (2 μM rotenone and 5 μM antimycin A, respectively) were carried out as indicated. The specific TRAP1 inhibitor compound 5 (100 μM) was added in fish water 4 hours prior to OCR analysis. In fish exposed to hypoxia, the drug was added 2 hours before hypoxic treatment and maintained throughout hypoxia exposure. The number of animals used for each condition is reported inside column bars. E Schematic representation of the feed-forward crosstalk between HIF1α and TRAP1. In (A-D), data are reported as average ± S.E.M. with one-way ANOVA and Bonferroni’s correction of 4 independent experiments (A, C, D), or at least 20 animals per condition (B) with an unpaired twotailed Student’s *t* test. Asterisks indicate significant differences (*p<0.05, **p < 0.01, ***p < 0.001).

## Discussion

In the present study, we demonstrate that the mitochondrial chaperone TRAP1 is a transcriptional target of HIF1α and that TRAP1 is induced in hypoxic conditions both in fish embryos and in tumor models. Under hypoxia, TRAP1 plays a major role in inhibiting respiration.

We have previously found that TRAP1 can prompt the HIF1α transcriptional program via SDH inhibition and the consequent raise in intracellular succinate that inhibits proteasomal degradation of HIF1α (Sciacovelli *et al*, 2013). These observations, together with those reported herein, indicate the presence of a feed-forward loop where TRAP1 induces HIF1α, which in turn increases TRAP1 levels (**Figure 5E**). This crosstalk could play important roles in coupling the metabolic status of the cell with environmental cues, such as fluctuating oxygen tension. These unstable conditions are experienced by cells not only in pathological conditions, exemplified by the growth of the neoplastic mass in an irregularly vascularized milieu (Vander Heiden & DeBerardinis, 2017, Ward & Thompson, 2012), but also in specific physiological settings. During embryonic development, conditions of variable oxygen usage frequently occur when cells must sustain high rates of biomass production during morphogenetic events. Indeed, developing embryos display high levels of aerobic glycolysis (Dunwoodie, 2009, Lange *et al*, 2016), highly similar to the Warburg effect that characterizes many cases of tumor growth, and HIF1α plays a master regulatory role in the correct differentiation of organs and tissues (Ietta *et al*, 2006).

TRAP1 is a critical regulator of mitochondrial metabolism under stress conditions (Masgras *et al*, 2017b, Rasola *et al*, 2014). Here we find that TRAP1 is highly expressed at the beginning of fish embryogenesis, when it exerts an important bioenergetic role. As previously reported in tumor models (Guzzo *et al*, 2014, Kowalik *et al*, 2016, Masgras *et al*, 2017a, Sanchez-Martin *et al*, 2020a, Sanchez-Martin *et al*, 2020b, Sciacovelli *et al*, 2013), TRAP1 inhibits SDH activity during early Zebrafish development, crucially contributing to a bioenergetic phenotype characterized by low levels of OXPHOS. Absence of TRAP1 causes a delay in early development; this defect is gradually lost during the passage from embryo to larva stages, when oxygen tension increases and TRAP1 levels progressively decline. These data indicate that TRAP1 is a component of the machinery that connects the efficient utilization of metabolism for developing purposes, albeit the lack of an overt phenotype caused by the homozygous deletion of TRAP1 in Zebrafish suggests that compensatory mechanisms must exist, in accord with what was already observed in TRAP1 knock out mice (Lisanti *et al*, 2014).

TRAP1 expression increases in both fish embryos and human cancer cells exposed to low oxygen tension. The TRAP1 promoter harbors HIF1α binding sites called hypoxia-responsive elements (HRE) in both Zebrafish and human genes, as well as binding sites for five additional transcription factors (SP1, HDAC1, SIN3A, MAX and MYC), which could cooperate with HIF1α in tuning TRAP1 expression. Interestingly, both SIN3A and MAX regulate the activity of MYC, and scattered evidences suggest that TRAP1 expression is elicited by MYC activity (Agarwal *et al*, 2019, Coller *et al*, 2000). Myc can cooperate with HIF1α to ensure the rapid adaptation to an environment endowed with oxygen paucity by modulating the expression of several target genes involved in metabolic reprogramming, cell cycle and proliferation (Cannino *et al*, 2018). Among HIF1α target genes, heat shock proteins (HSPs) are key during development (Hammerer-Lercher *et al*, 2001, Mestril *et al*, 1994), as they ensure correct protein folding and activity (Krone *et al*, 1997, Pechan, 1991) especially under stressful conditions such as hypoxia. The expression profile of the HSP90-family paralog TRAP1 well correlates with that of other HSPs modulated by HIF1α during Zebrafish embryogenesis (Elicker & Hutson, 2007).

Of note, TRAP1 induction occurs very rapidly also following pseudo-hypoxic stabilization of HIF1α. In cancer, this could be an adaptive mechanism to equip cells in advance for upcoming harsh conditions of nutrient and oxygen shortages. Blocking TRAP1 activity pharmacologically or by knocking-out its expression fully abrogates the inhibition of respiration that characterizes conditions of lower oxygen availability. Instead, un the presence of TRAP1, respiration is inhibited under hypoxia, and this downregulation is maintained even shortly after normal oxygen tension is restored. This finding position TRAP1 as one of the primary inhibitors of mitochondrial respiration under hypoxia. Its bioenergetic consequences could include protection of mitochondria from ROS produced during post-ischemic reperfusion and shaping metabolic responses that rely upon succinate accumulation.

The use of selective inhibitors of TRAP1 chaperone activity (Sanchez-Martin *et al*, 2020b) has proven instrumental here to demonstrate that these bioenergetic adaptations are rapidly reversible, which probably confers a high flexibility in harmonizing respiratory regulation with environmental changes. The chaperone activity of TRAP1 is enhanced by its ERK-mediated phosphorylation (Masgras *et al*, 2017a), and dysregulated activation of Ras/ERK signalling is mandatory for the malignant growth of a variety of cancer types, including highly aggressive pancreatic adenocarcinoma and malignant peripheral nerve sheath tumors, where we show that TRAP1 has a dramatic inhibitory effect on SDH activity (this study and (Sanchez-Martin *et al*, 2020b). Hence, the possibility of targeting TRAP1 with great precision unlocks the doorway to the development of therapeutic approaches in neoplastic conditions where the metabolic rewiring that it orchestrates is critical for malignant cell viability.

## Materials and Methods

### Animals

All Zebrafish studies were performed in accordance with the European and Italian legislation and the local ethical committee of University of Padova (authorization numbers 348/2020-PR and 239/2020-PR). Zebrafish embryos were maintained in a temperature-controlled room (28.5°C) and fed as described by Kimmel et al. (Kimmel *et al*, 1995). Fish were kept under a 14 h light-10 h dark cycle. For mating, males and females were separated in the late afternoon and the next morning were freed to start courtship, which ended with eggs deposition and fecundation. Eggs were collected, washed with fish water (0.5 mM NaH_2_PO_4_, 0.5 mM NaHPO_4_, 0.2 mg/l methylene blue, 3 mg/l instant ocean) and maintained at 28.5°C in fish water supplemented with an antibiotic-antimycotic cocktail (50 μg/ml ampicillin, 100 units/ml penicillin, 0.1 mg/ml streptomycin and 3.3 μg/ml amphotericin B). The male Zebrafish reporter line for HIF1α Tg(4xhre-tata:eGFP)^ia21^ (Vettori *et al*, 2017) was outcrossed with female wild type Zebrafish, and green heart positive offspring embryos were selected and used for experimental purposes. To generate a Zebrafish model of pancreatic adenocarcinoma, one cell stage eggs derived from Tg(ptf1a:Gal4) outcrossed with Wild Type Zebrafish were injected with Tol2(UAS:eGFP-KRAS^G12D^) plasmid, generating the final transgenic line Tg(ptf1a:Gal4)/UAS:eGFPKRAS^G12D^ (Schiavone *et al*, 2014). Approximately 100 embryos were raised, all expressing eGFP according to the expected ptf1a pattern. eGFP expression was evaluated from 1 week intervals to 1 month. Positive fish expressing eGFPKRAS^G12D^ were then monitored until development of a tumor mass, when fish were euthanized for further histological biochemical analysis. Fish were photographed live using a NIKON C2 H600L confocal microscope with 20× and 40× water dipping objectives. Laser used to excite fluorophores was 488 nm for eGFP. To generate Zebrafish TRAP1 knock-out model, eggs were injected at one cell stages and analysed at 24, 48 hpf and at 1 month post fertilization. The developmental stages at which experiments were carried out are indicated in the main figures and are within the 5^th^ dpf.

### Human cell lines

Experiments were performed on human U87 glioblastoma cells, human pancreatic carcinoma cells (MiaPaCa-2), human pancreatic adenocarcinoma cells (BxPc3) and human ipNF95.6 plexiform neurofibroma cells (kindly provided by Dr. Margaret R. Wallace, University of Florida, College of Medicine, Gainesville, FL). U87 cells were grown in Minimum Essential Medium (MEM) supplemented with 10% fetal bovine serum, 1% glutamine, 1% sodium pyruvate, 1% non-essential amino acids and 1% penicillin and streptomycin. MiaPaCa-2, BxPc3 and P.N. were grown respectively in Dulbecco’s Modified Eagle’s Medium (DMEM) and RPMI-1640 supplemented with 10% fetal bovine serum (FBS) 100 units/ml and 1% penicillin and streptomycin. All cells were cultured at 37°C in a humidified atmosphere containing 5% CO_2_

### sgRNA template design and generation

The CRISPR/Cas9 mediated mutagenesis of TRAP1 gene was achieved by applying the protocol proposed by Gnagnon at al. (Gagnon *et al*, 2014). CHOPCHOP (https://chopchop.cbu.uib.no/) was used to identify the sgRNA sequence against the exon 6 of TRAP1 gene. For the gene-specific oligonucleotide design, the SP6 promoter sequence was added before the sgRNA followed by an overlap sequence, which interacts with the 80 bp long constant oligo. The dsDNA was generated by using the T4 DNA polymerase (NEB M0203S) after annealing of gene specific oligonucleotide and the constant oligo. The dsDNA template was then purified by using the PCR clean up column (Quiagen 28704) and transcribed over night at 37°C with the Ambion Megascript SP6 KIT (Ambion AM1330). The resulting sgRNA was purified with RNA clean and concentrator Kit (Zymo Research R1017), quantified and stored at −80°C.

### sgRNA and Cas9 injection and determination of somatic mutagenesis rate

Zebrafish (Tübingen) eggs were collected and injected with a mix composed by sgRNA (70ng), Cas9 protein (NEB M0641S) and phenol red injection dye. The sgRNA and Cas9 were incubated 5 minutes at room temperature to form the complex. Embryos were injected soon after the fertilization and kept at 28.5°C. the day after injection the genomic DNA was extracted from 24 hpf injected embryos. Single fish were incubated 20 minutes with NaOH 50 mM at 95°C, then cooled on ice followed by addition of Tris-HCl 1 M, pH 7.5. The extracted DNA was amplified by PCR using Phusion High-fidelity Master mix (M0531S) and relative primers. We then performed heteroduplex mobility assay (HMA) to identify the presence of somatic mutations. Briefly, the PCR products were mixed with HMA buffer (1 M NaCl, 100 mM Tris (pH 7.8), and 20 mM EDTA) and gel loading buffer 1X. Samples were denatured at 95°C 15 minutes, then placed 15 minutes at room temperature and cooled on ice. Samples were loaded in a 10% polyacrilammide gel in TBE (boric acid 89 mM; EDTA 2 mM, pH 8.3) at 30 mA for 2 hours. Homoduplexes and heteroduplexes were visualized following staining of the gel with SYBR Green II nucleic acid stain (Invitrogen).

### Determination of germline mutations and generation of adult fish homozygotes

Upon sgRNA activity evaluation in somatic tissue, the remaining embryos were left to grow in order to identify germline-transmitting founders (F_0_). Each founder was outcrossed with wild-type Tübigen Zebrafish, and then 15 embryos from each pair were screened for mutations in F_1_. The DNA was extracted and amplified by PCR. Possible mutations were detected with HME on PCR products as described above. Although HMA is a rapid method to screen mutants, it does not provide the exact nature of the mutation. The screened alleles must be sequenced in order to get precise information on the mutation. Moreover, sequencing allows to discard those mutations that do not result in the introduction of a stop codon cassette. The positive F_1_ fish were raised to adulthood and crossed again with wild-type ones to generate the F_2_ population. Heterozygous were identified by PCR and sequencing of DNA extracted from tail at 1 month post fertilization. This scheme was used to generate F_3_ heterozygotes fish. Then, the F_3_ heterozygotes fish were inbred, and the homozygotes were identified by DNA extraction from tail at 1 mpf followed by PCR and sequencing analysis

### Generation of HIF1α knock-down cell lines

HIF1α knock-down cells were generated by using the short hairpin RNA strategy in U87 cell line. Sequences for the HIF1α shRNA (shRNA10819 5’-CCGGTGCTCTTTGTGGTTGGATCTACTCGAGTAGATCCAACCACAAAGAGCATTTTT-3’ and shRNA3809 5’-CCGGCCAGTTATGATTGTGAAGTTACTCGAGTAACTTCACAATCATAACTGGTTTTT-3’) were obtained from Sigma Aldrich. Scrambled shRNA was used as negative controls. shRNA oligonucleotide were cotransfected with the packaging plasmids psPAX (Addgene #12260) and pMD2.G (Addgene, #12259) into human embryonic kidney (HEK) 293T cells for viral production. Recombinant virus was collected and used to infect cells by standard methods. Infected cells were then selected with 2 μg/ml puromycin.

### Drug administration to fish and exposure to hypoxic conditions

Zebrafish embryos were maintained in hypoxia (5% O_2_) for the indicated time in a temperature-controlled room (28.5°C). Dimethyloxalylglycine (DMOG, 100 μM or 150 μM), dimethyl succinate (DMS, 1 mM) and TRAP1 inhibitor compound 5 (100 μM) were dissolved in fish water and maintained for the indicated time at 28.5°C. U87, MiaPaCa-2, BxPc3 and ipNF95.6 cells were incubated in hypoxia chamber (Baker Workstation InVIVO_2_ I400) at 0.5% O_2_, or treated with 0.5 mM CoCl_2_ or 100 μM TRAP1 inhibitor compound 5 for the indicated times.

### Epiboly, fish area and pigment measurements

To estimate epiboly and fish area the cell layer or the total fish body, respectively, were calculatedby excluding the yolk. For pigments analysis the trunk region was considered and the numbers of pigments were counted by using ImageJ software. All pictures were taken with Leica S9i microscope analyzed with ImageJ software. Fish area, epiboly and number of pigments are average ±SEM of the indicated embryos per condition.

### Protein isolation and Western Blot analysis

For Western immunoblot analysis, embryos or cells were lysed at 4°C in in RIPA buffer composed by Tris-HCl 50 mM pH7.4, NaCl 150 mM, NP40 1%, sodium deoxycholate 0.5%, SDS 0.1%, EDTA 2 mM and protease inhibitors (Sigma). Lysate were then clarified at 14000 rpm for 30 minutes at 4°C and quantified by using a BCA Protein Assay Kit (Thermo-Scientific). Protein extracted from embryos, tissue or cells were boiled at 50°C with Leamli buffer for 5 minutes, separated in reducing conditions by using NuPage Novex 4%-12% Bis-Tris gels (Life Technologies) and transferred in Hybond-C Extra membranes (Amersham). Primary antibodies were incubated for 16h at 4°C. Proteins were visualized using the UVITEC imaging system following incubation with horseradish peroxidase conjugated secondary antibodies.

### RNA extraction and RT-PCR

Total mRNA was isolated from Zebrafish embryos by using TRIZOL (Invitrogen). 2 μg of total mRNA was retro-transcribed with Superscript (Invitrogen). qPCR was performed by using GoTaq qPCR Master Mix (Promega).

### Measurement of succinate:coenzyme Q reductase (SQR) activity of SDH

To measure the succinate dehydrogenase (SDH) activity embryos were homogenized in an assay buffer composed by KH_2_PO_4_ (25 mM, pH 7.4) and MgCl_2_ (5 mM) containing protease and phosphatase inhibitors (Sigma Aldrich). Total lysates were then quantified using a BCA Protein Assay Kit (Thermo-Scientific). 60 μg of total homogenate was incubated for 10 minutes at 30°C in the assay buffer supplemented with sodium succinate 20 mM and alamethicin 5 μM. After incubation, a mix composed by sodium azide 5 mM, Antimycin A 5 μM, Rotenone 2 μM and Coenzyme Q1 65 μM was added. SDH activity was measured by following the reduction of 2-6 dichloro phenolindophenol (DCPIP) at 600 nm (ε=19.1 nM^−1^ cm^−1^) at 30°C. Inhibitors (17AAG or compound 5) were added 5 minutes before starting recordings.

SDH enzymatic activity was also assessed histochemically on frozen tissue. Soon after extraction, pancreatic tissue were embedded in frozen OCT (Kaltek 0782). The tissue sections were incubated 30 minutes at 37 °C with incubation medium containing KH_2_PO_4_ 0.2 M, pH 7.4, sodium succinate 0.2 M and nitro blue tetrazolium (NBT) salt (N6876 Sigma-Aldrich). The TRAP1 inhibitor compound 5 (100 μM) was included in incubation media were indicated. Sections were then rinsed with PBS 1X and covered by using a mounting medium. Succinate was oxidized to fumarate and reduction of the final electron acceptor nitro blue tetrazolium (NBT) allows the formation of a purple staining.

### Measurement of oxygen consumption rate (OCR)

Oxygen consumption rate was measured with Seahorse XF24 extracellular flux analyzer in living Zebrafish embryos at 96 hours post fertilization (hpf). Zebrafish embryos were placed into XF24 microplate well (1 embryos per well) and blocked with a capture screen to keep them in place; 670 μl of fish water (0.5 mM NaH_2_PO_4_, 0.5 mM NaHPO_4_, 3 mg/l instant ocean) was added. The basal respiration was measured for 130 or 68 minutes at 28.5 °C. FCCP (0.5 μM) was used to measure maximal respiration while Rotenone (0.5 μM) and Antimycin A (5 μM) were used to completely abolish mitochondrial respiration. Respiratory rates are average±SEM of the indicated embryos per condition.

### *In silico* analysis of TRAP1 promoter

The TRAP1 nucleotide sequences were retrieved from Ensembl (Aken *et al*, 2016) (accession codes: ENSG00000126602 and ENSDARG00000024317 for human and zebrafish, respectively), aligned with BLASTn (Zhang *et al*, 2000) and visualized with Jalview (Waterhouse *et al*, 2009). Conserved HREs were extracted through bash shell command line, while transcription factors binding within the human TRAP1 promoter region were retrieved from GeneCards (Stelzer *et al*, 2011). HIF-1α interactors were extracted from the STRING (Szklarczyk *et al*, 2017).

### *In vivo* measurements of ROS levels

ROS levels were measured by incubating WT and TRAP1 KO Zebrafish larvae at 96 hpf in Hank’s Balanced Salt Solution (HBSS) and MitoSOX probe for 30 minutes at 30°C. Then, three wash of ten minutes were performed. Red fluorescence emitted by mitoSOX probe was measured by acquiring live embryos image with a Nikon C2 H600L confocal equipped with 40X water immersion objective, camera and laser emitted at 561 nm. ROS signal was analyzed by measuring the integrated density of emitted fluorescence in the brain region normalized for the area analyzed with ImageJ.

### Antibodies

Mouse monoclonal anti-rodent TRAP1 was from Becton Dickinson (Cat. # 612344); mouse monoclonal anti-human TRAP1 was from Santa Cruz (Cat. # sc-73604); mouse monoclonal anti-β actin was from Santa Cruz (Cat. # sc-47778); rabbit polyclonal anti-citrate synthetase was from Abcam (Cat. # ab96600); rabbit polyclonal anti-HIF1α was from Novus Biologicals (Cat. # NB100-449); rabbit polyclonal anti-HIF1α was from GeneTex (Cat. # GTX131826); rabbit polyclonal anti-phospho-ERK1/2 was from Cell Signaling (Cat.# 9101); rabbit polyclonal anti-glucose transporter GLUT1 was from Abcam (Cat.# ab652).

### Quantification and statistical analysis

Data were analyzed and presented as mean ± standard error of the mean (SEM) in all figures. Pairs of data groups were analyzed using paired and unpaired two-tailed Student’s *t* tests. In the case of more than two groups, one-way analysis of variance (ANOVA) followed by Bonferroni post hoc test was applied. Statistical significance was determined using GraphPad Prism 8. Results with a p value lower than 0.05 were considered significant; ***p < 0.001, **p < 0.01, *p < 0.05 compared to controls. Each experiment was repeated at least three times.

## Acknowledgments

A.R. was supported by grants from University of Padova, Neurofibromatosis Therapeutic Acceleration Program and Associazione Italiana Ricerca Cancro (AIRC grant IG 2017/20749). G.C was supported by a grant from Associazione Italiana Ricerca Cancro (AIRC grant IG 2017/20019). I.M. was recipient of a Young Investigator Award Grant from Children’s Tumor Foundation. We thank the Busch-Nentwich lab for providing RNA-seq data and Elena Trevisan for excellent technical assistance.

## Author contributions

Conceptualization: C.L., I.M. and A.R.; visualization, C.L.; methodology: C.L., M.S., S.T., F.A., G.C., P.B., I.M., and A.R.; investigation: C.L., C.S.M., G.M., E.M., I.M.; formal analysis: C.L., C.S.M., G.M., E.M., G.C., I.M. and A.R.; resources: F.A., S.T. and P.B.; writing-original draft: C.L., I.M. and A.R.; writing-review and editing: C.L., P.B., I.M. and A.R.; funding acquisition; A.R.; project administration: A.R.; supervision: I.M. and A.R.

---

**Table 1.**
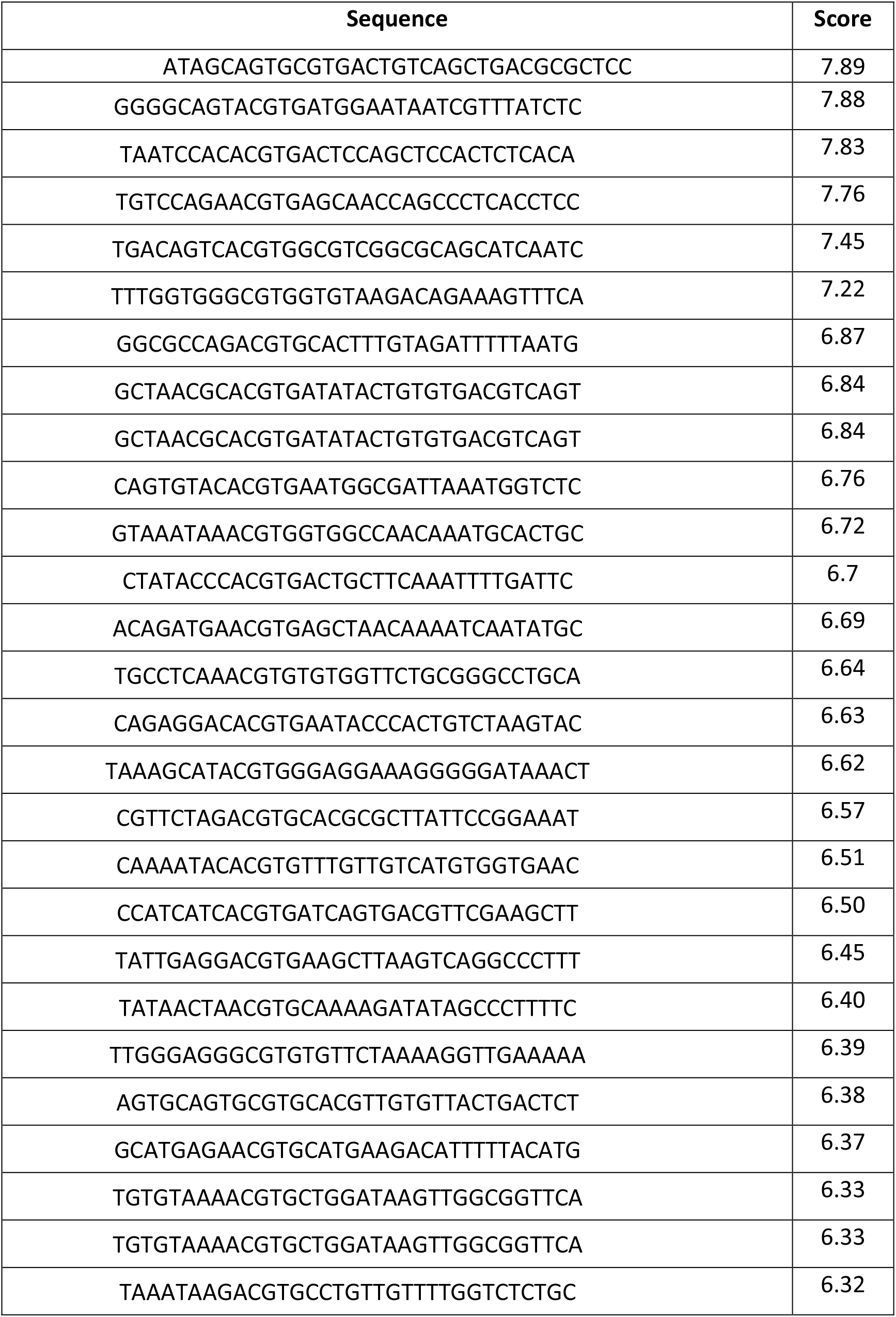
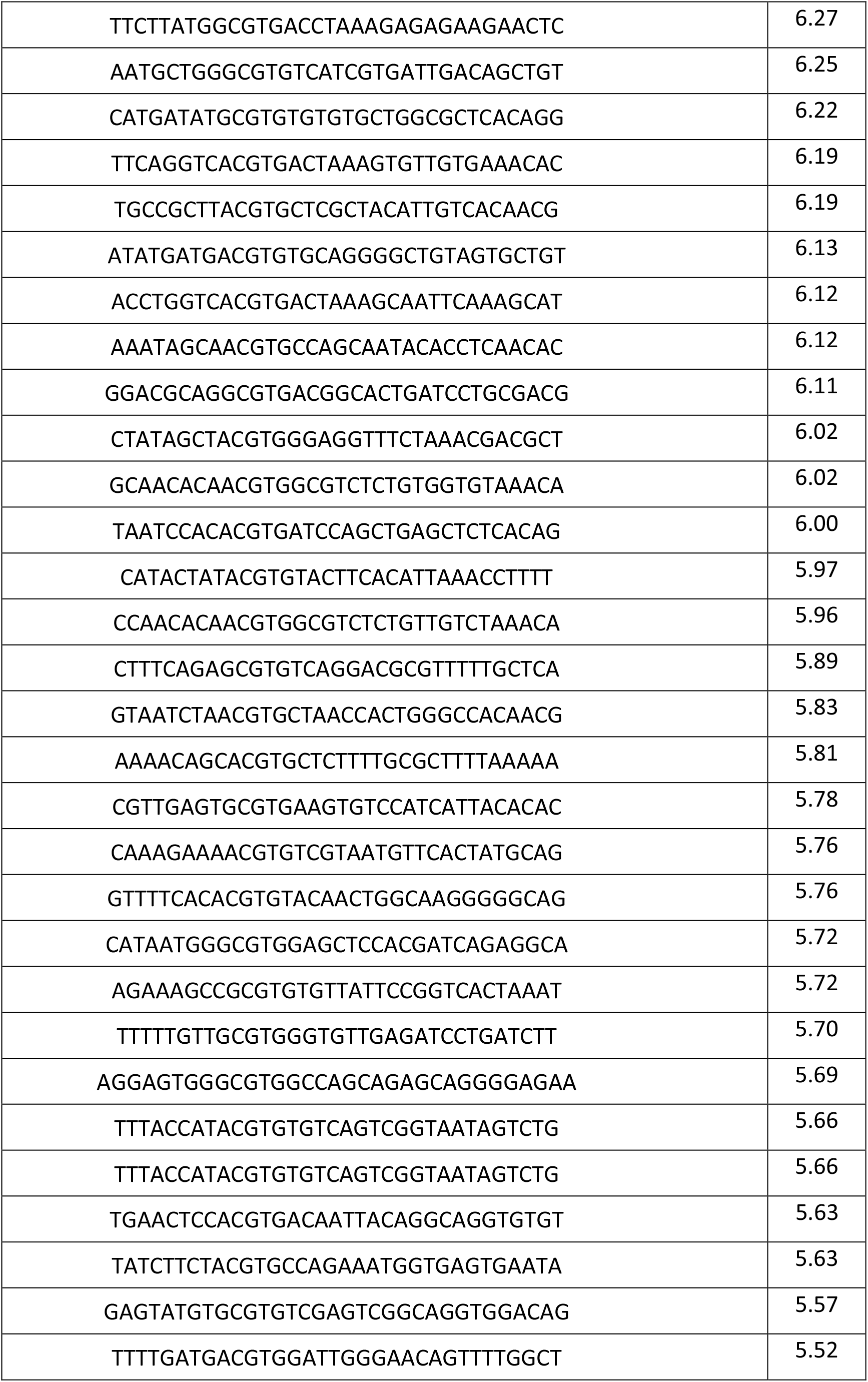
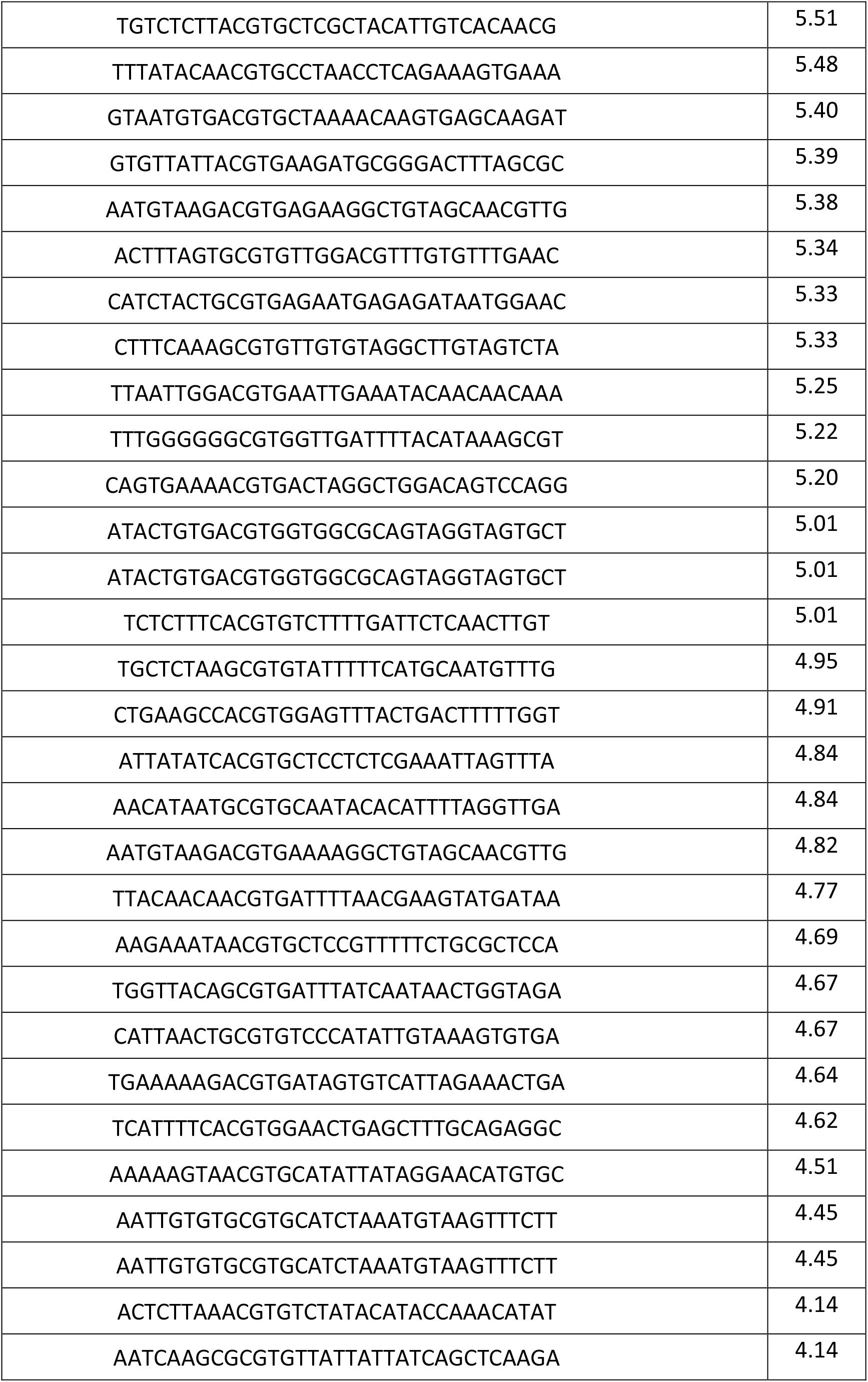
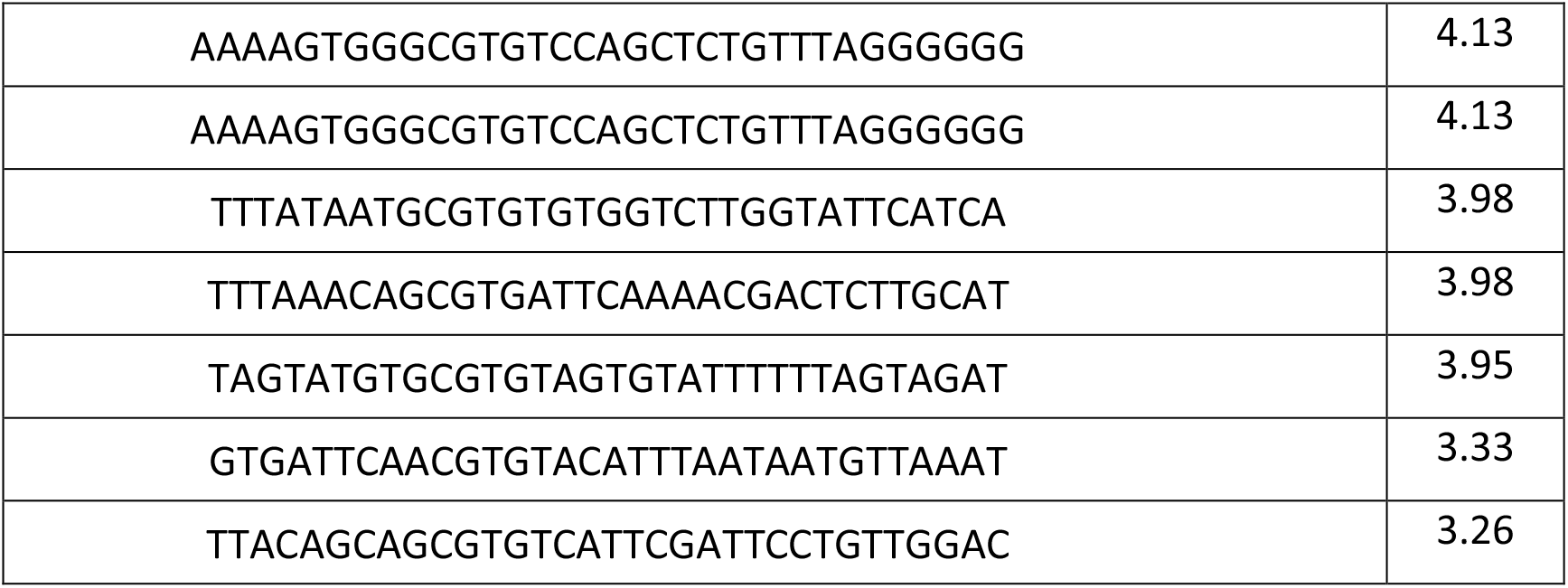
94 positive sequences obtained in the preliminary screening of Zebrafish TRAP1 promoter. The sequences are homogeneously distributed, showing no preference of localization.

**Table 2.**
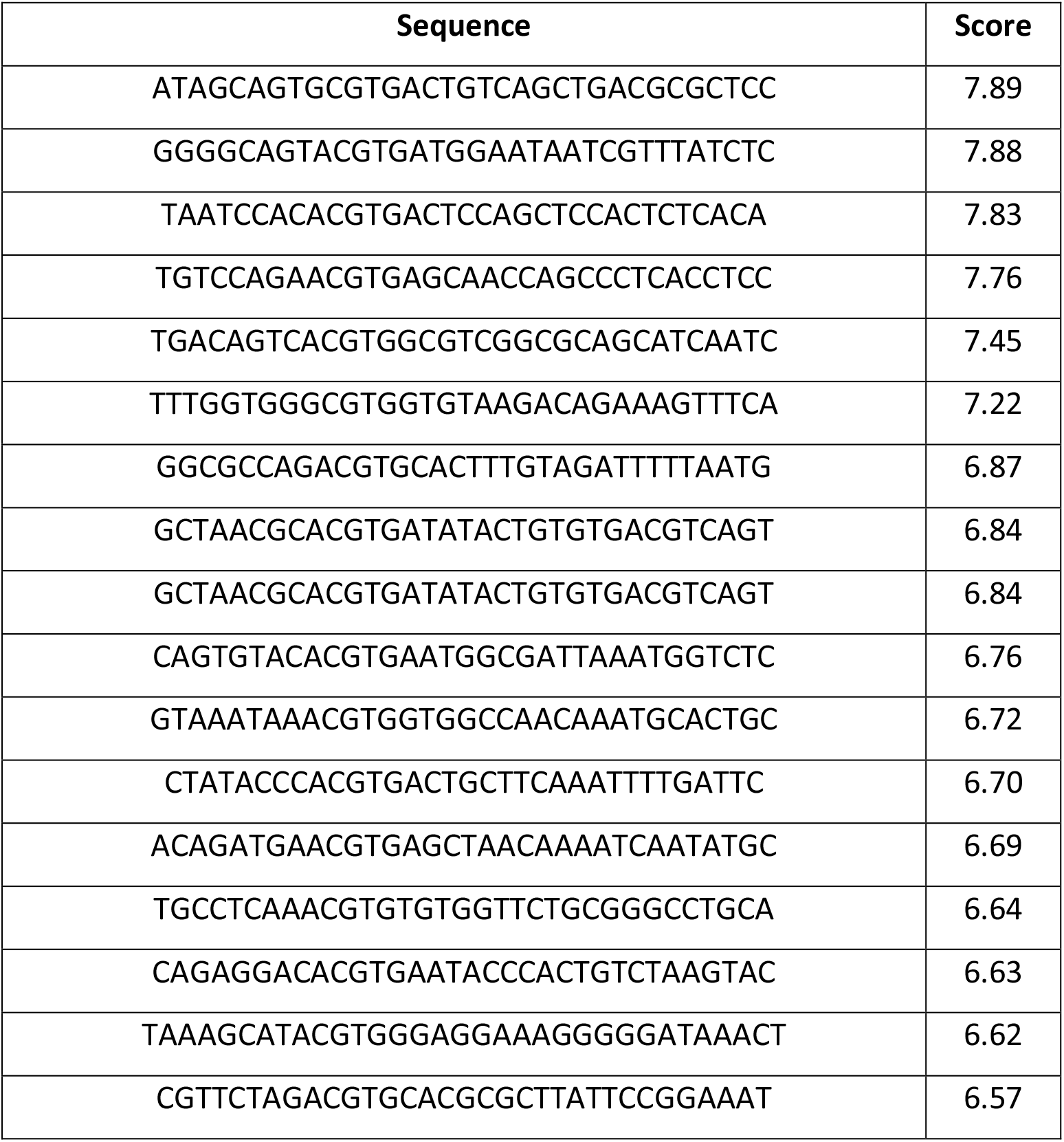
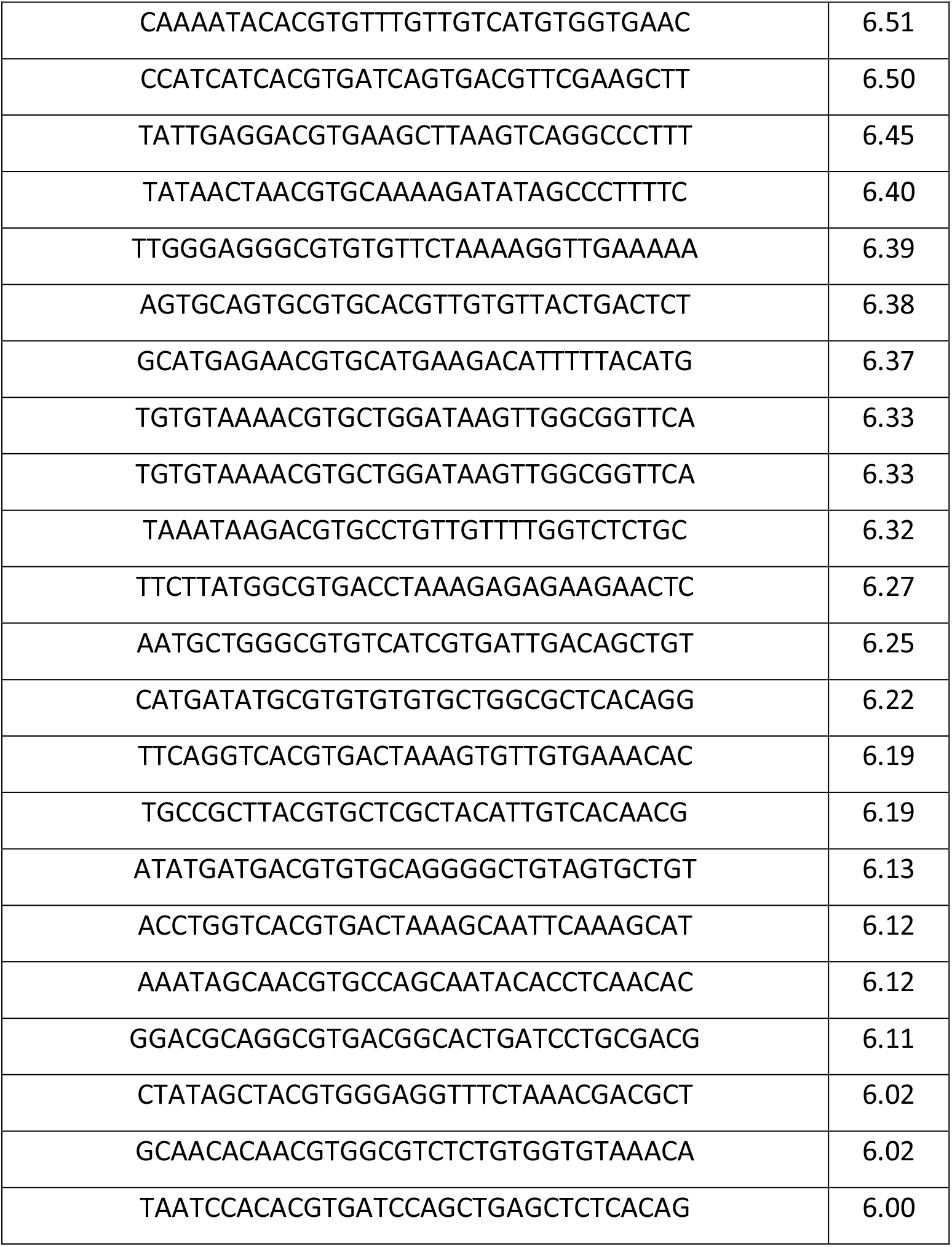
List of the top ranking HREs identified within position −5000bp and +500bp from the zebrafish TRAP1 transcription initiation site (Dario rerio, Chromosome 3: 9,602,709-9,659,449) with associated probability score.

**Table 3.**
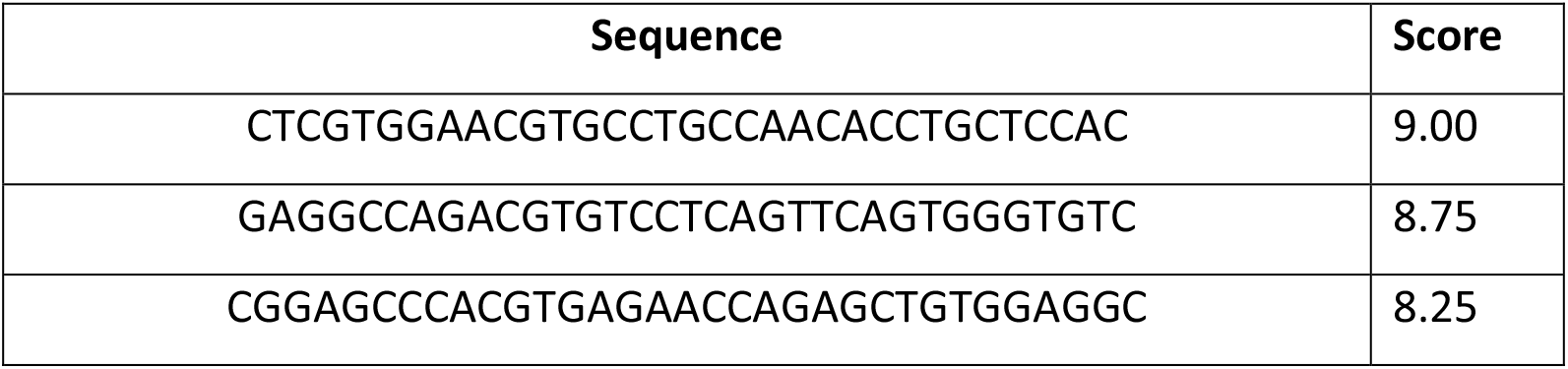
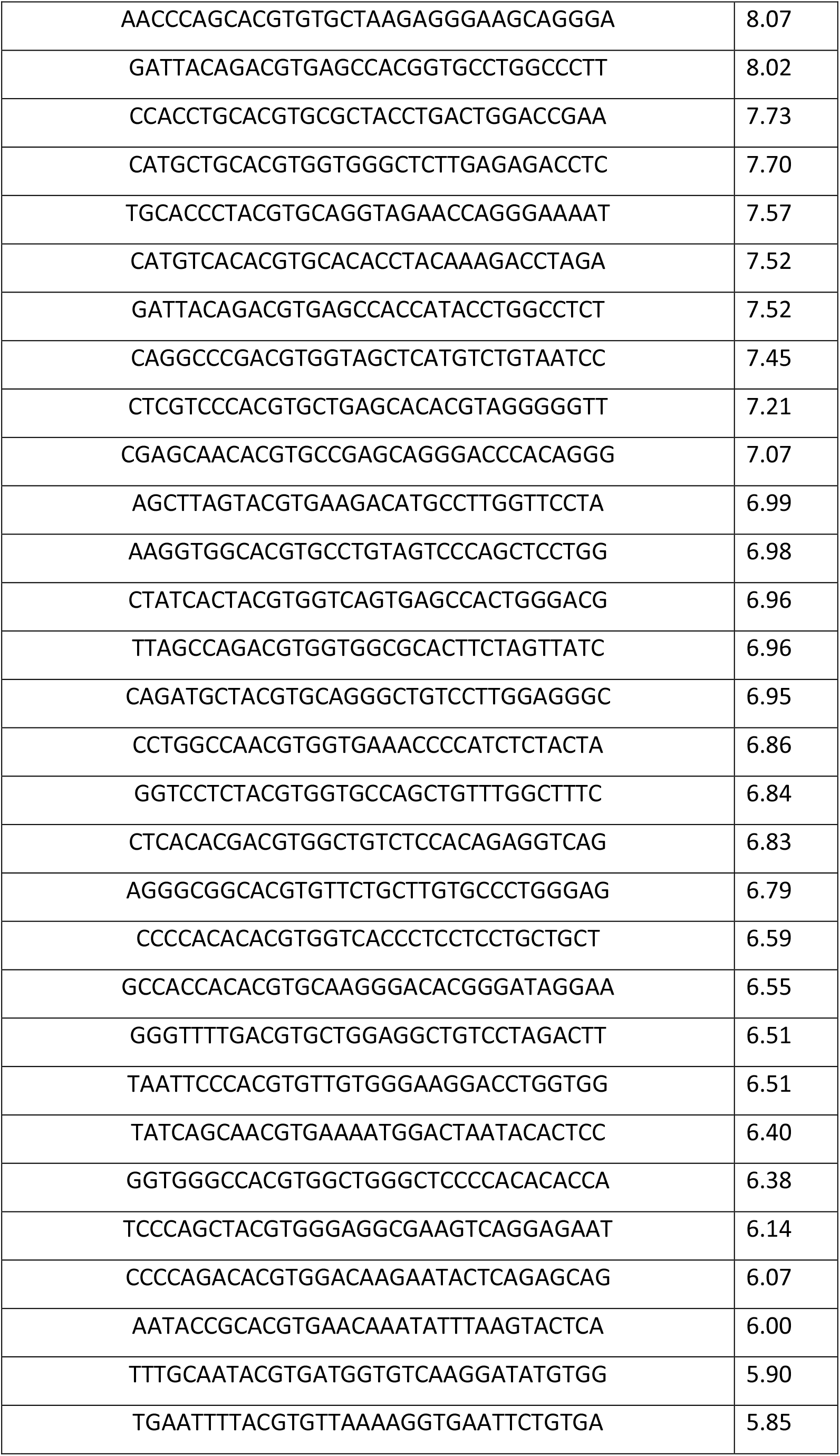
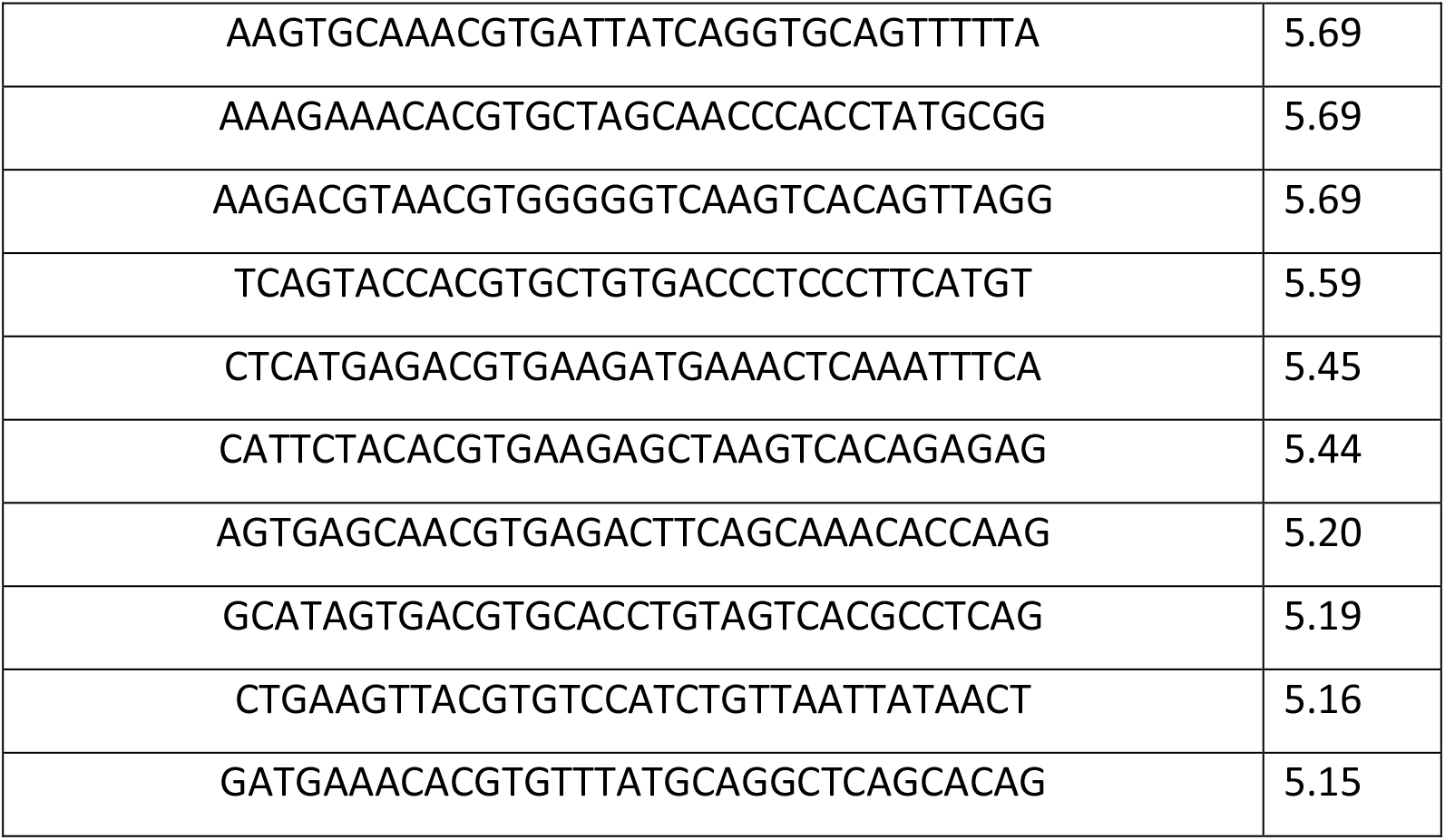
List of the top ranking HREs identified within the human TRAP1 promoter region (Homo sapiens, Chromosome 16: 3,716,198-3,718,509) with associated probability score.

**Table 4.**
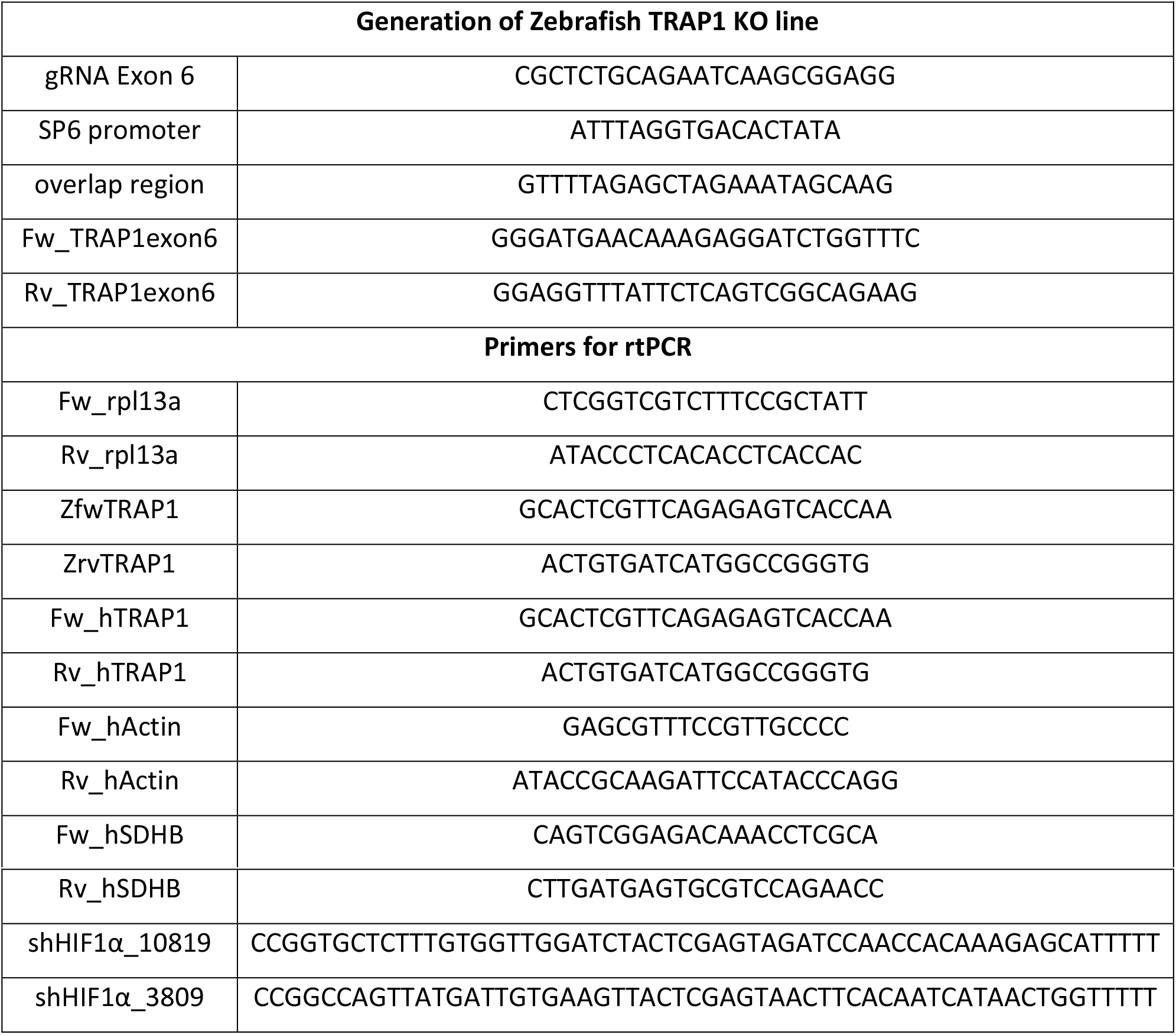
List of primers used to generate and to screen the Zebrafish TRAP1 knock-out fish, and primers used for real-time PCR

## Expanded view figure legends

**Figure EV1. Related to Figure 1. Generation of the TRAP1 knock-out fish**

A Schematic representation of the crossbreds to generate the final homozygous TRAP1 mutant genotype.

B Heteroduplex Mobility Assay (HMA) on PCR products of total DNA extracted from F1 Zebrafish indicates the presence of heterozygous mutation in lane #2.

C PCR- and sequencing-based screenings identify the presence of homozygous mutation in TRAP1 exon 6 (deletion of 13 nucleotides, Δ13). Genotyping PCR identifies the presence of 275 bp and 262 bp PCR product in wild type and mutated TRAP1 allele, respectively.

D Amplification of TRAP1 cDNA.

E Western blot analysis of TRAP1 protein level in wild-type and knock-out animals at 1 mpf.

F Heatmap and boxplot showing the relative TRAP1 gene expression level (TPM, transcription per million) across all 18 Zebrafish developmental stages from Expression Atlas. Five biological replicates were used for each developmental stage (http://www.ebi.ac.uk/gxa/experiments/E-ERAD-475).

G Measurement of mitochondrial ROS (Mitosox fluorescence) in TRAP1 wild-type and knock-out fish. Each fluorescent signal was normalized to the area of the analyzed region (head). Data are reported as average ± S.E.M. with an unpaired two-tailed Student’s *t* test of 6 animal per condition.

H Expression of HIF1α Tg(4Xhre-tata:GFP)ia21 Zebrafish reporter line during the first 5 days of Zebrafish development; left, quantification of integrated density fluorescence emitted by the HIF1α reporter line; right, representative images. Each fluorescent signal was normalized to the area of the analyzed region (head). Data are reported as average ± S.E.M. with one-way ANOVA and Bonferroni’s correction of at least 10 embryos per condition. Asterisks indicate significant differences (*p<0.05, **p<0.01, ***p<0.001).

**Figure EV2. Related to Figure 2. TRAP1 is induced by hypoxia in human tumor cell models**

A, B, C Western blot analysis of TRAP1 expression profile in human glioblastoma U87 cells (A), pancreatic adenocarcinoma BxPc-3 cells (B) and human ipNF95.6 plexiform neurofibroma cells (C) cell lines upon hypoxia exposure for the indicated time. CoCl2 (0.5 mM for 6 hours) was used as a positive control for HIF1⍰ stabilization.

**Figure EV3. Related to Figure 3. TRAP1 is regulated in a HIF1α-dependent manner**

A Putative network of TRAP1 transcriptional regulators. Transcription factors known to bind HIF1α and that bind the TRAP1 promoter region according to inspection with STRING database were connected. Magenta edge lines represent direct interactions, while purple lines are used for co-expression relationships.

B Model of TRAP1 gene transcriptional regulation derived from the above in silico analysis. HRE, hypoxia responsive element; TFs, transcription factors.

C Western blot analysis of TRAP1 expression in human U87 glioblastoma cells following HIF1α silencing and exposure to hypoxia. Data are reported as average ± S.E.M of at least 3 independent experiments with an unpaired two-tailed Student’s *t* test; asterisks indicate significant differences (*p<0.05, **p<0.01, ***p<0.001).

**Figure EV4. Related to Figure 4. Analysis of Zebrafish pancreatic tumors**

A Confocal images of pancreas at 1 mpf in ptf1a:Gal4/UAS:eGFP;Tg(4xHRE-tata:mcherry-nls) transgenic used as control (upper image) and in ptf1a:Gal4/UAS:eGFP-KRASG12D;Tg(4xHRE-tata:mcherry-nls) transgenic fish (lower image). In control fish the eGFP is expressed in the cytoplasm, staining the entire tissue, while the mcherry signal that indicates HIF1α activity is not present. In the transgenic model, eGFP-KrasG12D expression is strictly associated with plasma-membrane and the mcherry-nls signal is localized in the nucleus.

B Histologic analysis of adult fish pancreas showing malignant properties with mixed acinar and ductal features in the transgenic model (right); on the left, pancreatic tissue of wild-type animals.

C TRAP1 expression profile in Zebrafish tissues at 1 ypf.

**Figure EV5. Related to Figure 5. Effects of TRAP1 inhibition on hypoxic Zebrafish embryos**

A Measurement of fish area in TRAP1 wild-type and knock-out fish kept in normoxic or hypoxic (5% O_2_) conditions for 72 hours (from 6 hpf up to 72 hpf).

B Analysis of succinate dehydrogenase (SDH) activity on wild-type Zebrafish embryos kept in normoxia or hypoxia for 48 hours. Where indicated, embryos were treated with the wide-spectrum HSP90 family inhibitor 17-N-allylamino-17-demethoxygeldanamycin (17AAG, 10 μM).

Data are reported as average ± S.E.M. of at least 15 animals per condition with an unpaired two-tailed Student’s *t* test. Asterisks indicate significant differences (*p<0.05, **p<0.01, ***p<0.001).

